# A humanness dimension to visual object coding in the brain

**DOI:** 10.1101/648998

**Authors:** Erika W. Contini, Erin Goddard, Tijl Grootswagers, Mark Williams, Thomas Carlson

## Abstract

Neuroimaging studies investigating human object recognition have largely focused on a relatively small number of object categories, in particular, faces, bodies, scenes, and vehicles. More recent studies have taken a broader focus, investigating hypothesised dichotomies, for example animate versus inanimate, and continuous feature dimensions, such as biologically similarity. These studies typically have used stimuli that are clearly identified as animate or inanimate, neglecting objects that may not fit into this dichotomy. We generated a novel stimulus set including standard objects and objects that blur the animate-inanimate dichotomy, for example robots and toy animals. We used MEG time-series decoding to study the brain’s emerging representation of these objects. Our analysis examined contemporary models of object coding such as dichotomous animacy, as well as several new higher order models that take into account an object’s capacity for agency (i.e. its ability to move voluntarily) and capacity to experience the world. We show that early brain responses are best accounted for by low-level visual similarity of the objects; and shortly thereafter, higher order models of agency/experience best explained the brain’s representation of the stimuli. Strikingly, a model of human-similarity provided the best account for the brain’s representation after an initial perceptual processing phase. Our findings provide evidence for a new dimension of object coding in the human brain – one that has a “human-centric” focus.

## Introduction

Human object recognition is fast, efficient (Thorpe, Fize, & Marlot, 1996) – and fundamental to our interactions with the world. The ventral temporal cortex (VTC) is widely accepted as a key structure for visual object perception (Caramazza & Shelton, 1998; Haxby, et al., 2001; Ishai, Ungerleider, Martin, Schouten, & Haxby, 1999; Mahon, et al., 2007). One hypothesized organisational principal in human and primate VTC the animate-inanimate dichotomy (Kiani, Esteky, Mirpour, & Tanaka, 2007; Kriegeskorte, Mur, Ruff, et al., 2008; Pinsk, et al., 2009). In support of this view, neuroimaging studies have shown subregions of the VTC with distinct response preferences, including a medial to lateral organization of animate and inanimate objects in the brain (Chao, Haxby, & Martin, 1999; Kanwisher, McDermott, & Chun, 1997; Konkle & Caramazza, 2013; Mahon, et al., 2007; Taylor & Downing, 2011). It is also well known that specific regions within VTC respond preferentially to images from particular categories, including faces, animals, bodies (Downing, Chan, Peelen, Dodds, & Kanwisher, 2006; Downing, Jiang, Shuman, & Kanwisher, 2001; Haxby, et al., 1994; Puce, Allison, Asgari, Gore, & McCarthy, 1996; Sergent, Ohta, & MacDonald, 1992), tools (Chao, et al., 1999; Chao & Martin, 2000) and places (Epstein, Harris, Stanley, & Kanwisher, 1999; Epstein & Kanwisher, 1998; Taylor & Downing, 2011). These category distinctions, notably however, represent a small sample of the wide array of objects that we see in everyday life.

An alternative approach to understanding object representations in the brain is to study how objects are coded in distributed patterns of brain activity (Haxby, et al., 2001; Ishai, et al., 1999) Using multivariate pattern analysis (MVPA) (for review see Grootswagers, Wardle, & Carlson, 2017; Haynes, 2015; Pereira, Mitchell, & Botvinick, 2009), researchers can study patterns of brain activity and test hypotheses about the neural representation of object categories (Kriegeskorte & Kievit, 2013; Kriegeskorte, Mur, & Bandettini, 2008). Using the MVPA framework, studies examining the relative similarity/dissimilarity of individual object representations in VTC’s have evidenced that objects may be represented along continuous dimensions in a multidimensional representation space. Animate subcategories have been argued to be coded along an axis of biologically similarity to humans (Connolly, et al., 2012; Sha, et al., 2015). This animacy continuum, however, does not provide a clear prediction for subcategory differentiation within the inanimate domain, nor for how the brain would represent objects that blur the animate-inanimate distinction (e.g., robots and animal toys). Moreover, it is also unclear whether a continuum centred around ‘animacy’ best captures the dimension along which neural responses vary. Sha et al. (2015), for example, proposed that the neural representation of objects is better characterised according to the object’s ability to perform goal-directed actions (see also Thorat, Proklova, & Peelen, 2019). Critically, there are many related factors to biologically similarity and agency that are known to influence human perception of objects (Gobbini, et al., 2011; Gray, Gray, & Wegner, 2007). This raise the question about whether these factors also might be used as organisational principles for the brain’s representation of objects.

In the present study, we used magneto-encephalography (MEG) to characterise the brain’s neural representations of objects, and to explore their temporal dynamics. We studied the brain’s emerging representation of 120 object stimuli and tested a wide range of models that might account for the brain’s representation of these objects using the representational similarity analysis (RSA) framework (Kriegeskorte & Kievit, 2013; Kriegeskorte, Mur, & Bandettini, 2008). We found that, after an initial period of perceptual processing, higher order category models and models of agency and human-related experiences account for brain’s representations of objects. Notably, the model that best accounted for later stage representations of objects was a “human-centric” model, which describes objects in terms of their similarity to humans.

## Materials and Methods

### Participants

Twenty-four English-speaking volunteers (18 female) with an average age of 24.7 years (SD = 5.47; range = 18-37) were recruited from the Macquarie University community. Informed written consent was obtained prior to participation, and participants were financially compensated for their time. All participants self-reported normal or corrected-to-normal vision (wearing of contacts was allowed), were free of medical conditions, and were not currently taking any neuroactive medications. This study was approved by the Macquarie University Human Research Ethics Committee.

### Stimuli

Stimuli consisted of 120 naturalistic images of objects (Figure 1), which were displayed on a uniform grey background. Twelve object categories were used in the study: six animate (humans, primates, domestic animals, birds, fish, invertebrates) and six inanimate (plants, robots, machines, tools, toys, other non-moving objects). In this stimulus set, animate is defined as living animals, in line with previous research (Caramazza & Shelton, 1998; Carlson, Tovar, Alink, & Kriegeskorte, 2013; Connolly, et al., 2012; Gobbini, et al., 2011; Kriegeskorte, Mur, Ruff, et al., 2008; Sha, et al., 2015). Categories were selected to include ones similar to those used by Sha et al. (2015), with the addition of robots and toys to address the questions about agency and experience. We also included machines, which, like robots, had moving parts, but did not have the humanistic/animalistic/agentic properties. Stationary objects were also included, which neither moved nor had humanistic/animalistic/agentic properties.

**Figure 1.**
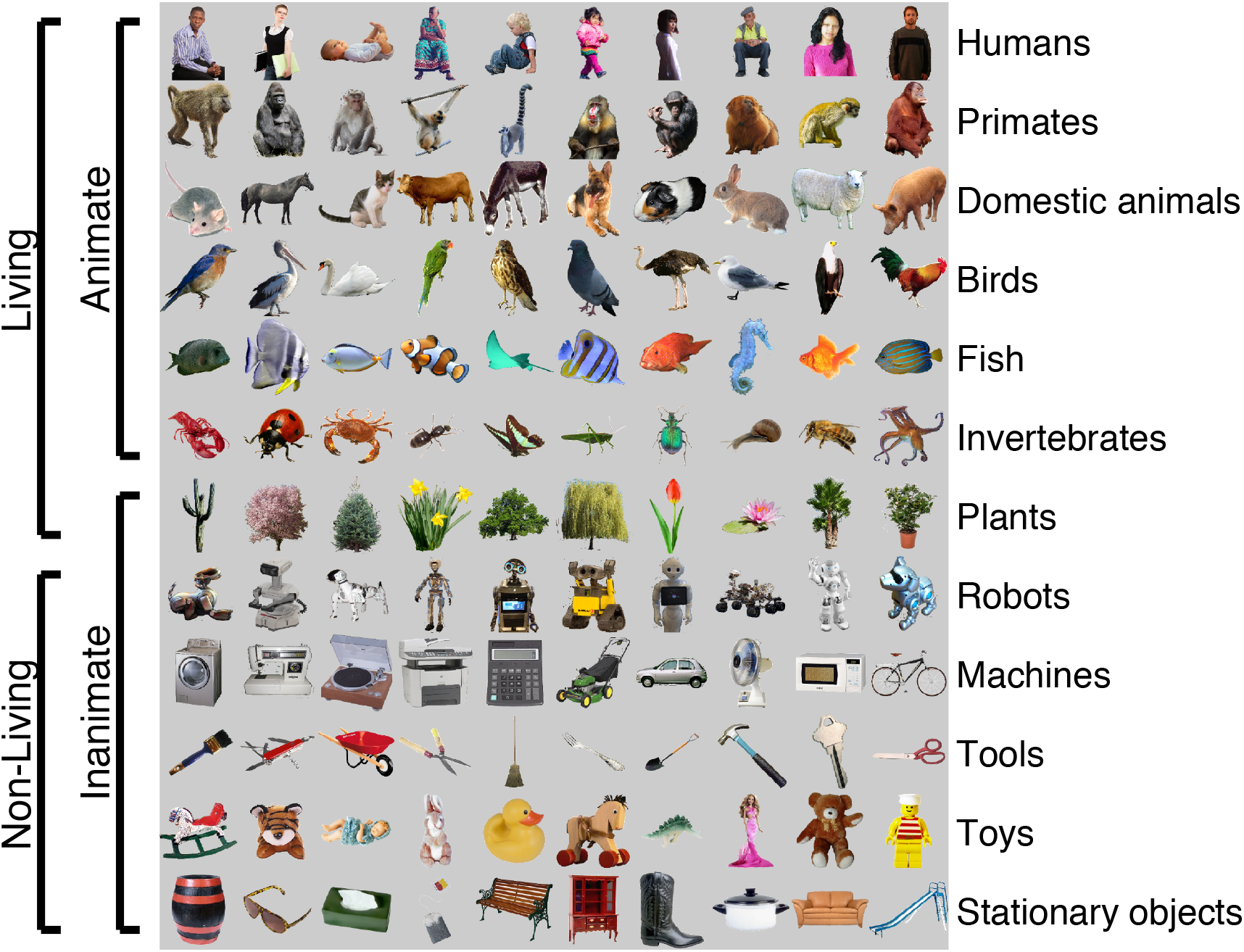
Stimuli from each of the 12 object categories. Animate object categories are ordered vertically according to the biological classes animacy continuum (Sha et al., 2015). Brackets show two examples of different groupings of the stimuli: living vs. non-living and animate vs. inanimate.

### MEG Experimental Procedure

For the experimental task, participants completed eight blocks of 398 trials (3184 trials in total). Within each block exemplars were presented for 100 ms, with a random inter-trial interval ranging between 750 and 1000 ms. The eight blocks were collected in a single session totalling approximately one hour of MEG recording time. Stimuli were presented in a predetermined pseudo-randomised order, such that for each trial, the preceding and following images had an equal probability of being from any one of the 12 object categories. The ordering of the 8 blocks was pseudo-randomised across participants.

Across trials, object images were manipulated in two ways to reduce the effects of low-level stimulus properties on our data. Firstly, a left-right flipped version of each image was included in the stimulus set, resulting in a total of 240 stimuli from 120 object images. Secondly, during image presentation, stimuli appeared in one of four locations while participants maintained fixation on a central marker, thus varying retinal location of the stimulus images. The four locations were defined by a shift from central presentation towards each of the four corners of the screen, where each stimulus location overlapped the central fixation point (details in *Display Apparatus* below). Each stimulus was presented three times at each location. This resulted in a total of 2880 trials (240 stimuli x 4 locations x 3 repetitions = 2880 trials). The additional trials were not included in the analysis: these included the first and last trial of each block, as well as 288 repeat trials that were added for the attention task (see below).

### Attention Task

During the experiment, participants completed a one-back attention task, where they were required to press a button whenever an object image was repeated consecutively. Participants received feedback about their accuracy on the task at the completion of each block. The mean accuracy across participants was 87.38% (*SD* = 7.28%), with an average reaction time of 535 ms (*SD* = 51 ms). Due to a malfunction of the response button during the experiment, accuracy and reaction times were missing for one of our 24 participants, as well as for one out of the eight blocks for each of two further participants. These participants were still instructed to perform the task and were unaware that the button was not recording their responses.

### Display Apparatus

Participants lay supine in the magnetically shielded recording room. Using an InFocus IN5108 projector situated outside the chamber, stimuli were projected onto a mirror, which reflected the image onto the ceiling, located approximately 113 cm above the participant. The total screen area was 20×15 degrees of visual angle (DVA). Throughout the experiment the screen background was held at a mean grey, and subjects were instructed to fixate on a black central fixation point (diameter of 0.1 DVA) that was always present. All stimulus locations were within a 6.9 DVA square, centred on the fixation point. Each stimulus consisted of a 256×256 pixel image (containing the segmented colour object) that was drawn to a 4.9×4.9 DVA square. Stimuli were presented one at a time, in one of four locations aligned with the upper left, upper right, lower left, or lower right corner of the 6.9 DVA square. A central square of 150 pixels (2.9 DVA) was common to all four stimulus locations. All stimuli were drawn as full colour segmented objects against a mean grey background (as in Figure 1): the same mean grey as the screen outside the stimulus location. Upon stimulus presentation, a 50×50 pixel (1×1 DVA) white square simultaneously appeared in the bottom right corner of the projection, which was aligned with a photodetector attached to the mirror to accurately record the stimulus presentation time in the MEG recording. The experiment was run on a Dell PC desktop computer using MATLAB software (Natick, MA) and the Psychophysics Toolbox extensions (Brainard, 1997; Kleiner, Brainard, & Pelli, 2007; Pelli, 1997).

#### MEG Data Acquisition

MEG data were recorded in the KIT-Macquarie Brain Research Laboratory using a 160-channel whole-head axial gradiometer (KIT, Kanazawa, Japan). Continuous data were acquired at a sampling rate of 1000 Hz, and were band-pass-filtered online from 0.03 to 200 Hz. MATLAB (2013b, Natick, MA) was used for all processing and statistical analyses of the data. Offline, we down-sampled the data to 200Hz and epoched each trial into an event with a time window from −100 ms to 600 ms relative to stimulus onset. To reduce the dimensionality of the data, we applied Principal Components Analysis to the epoched data from the 160 gradiometers, and retained the first *n* components that accounted for 99% of the variance. The number of components retained for each participant ranged from 14 to 72 (Mean = 34.21, SD = 18.90).

#### Classification analysis

For each participant, we used linear discriminant analysis to classify object/exemplar identity at the single trial level, training and testing classifiers on their ability to discriminate every possible exemplar pair of the 120 object images. We used cross-validated classification accuracy as a measure of how dissimilar the patterns of brain activity were for one exemplar compared to another (Nili, et al., 2014). We did not attempt to model the effects of spatial position or left-right flip in our classification analysis, but instead used a single data label (the object identity) for data obtained from both the standard and left-right-flipped versions of the stimuli, as well as all four stimulus presentation locations. By including data from all variations of the stimuli, we sought to force the classifier to generalise beyond lower-level visual features, (such as the presence or absence of stimulation at a given location in the visual field), and instead use any neural correlate of object identity. These modifications to the stimulus presentation would have introduced extra noise into the signal across trials, so would tend to reduce classifier performance relative to unvarying stimuli, but they allowed us to better target higher-level object representations. For each time-point, we trained and tested a separate classifier to discriminate each pair of exemplar identities from the PCA components. We used a 10-fold cross-validation procedure, where the classifier was trained on data from 90% of the trials and then its accuracy was evaluated using its performance when classifying the remaining 10% of the data, so that the classifier was never tested on data that were included in the training set. This process was repeated 10 times, so that all trials were used as test data once each. D-prime (d’) was used as the metric for classification accuracy.

#### Representational Similarity Analysis (RSA)

Classifier accuracies (d’) were averaged across exemplar pairs to obtain the mean classifier performance for each time point. Additionally, to capture the pattern of classifier performance across exemplar pairs and compare this pattern with model predictions, we constructed a Representational Dissimilarity Matrix (RDM) for each time point. The RDM is a 120×120 matrix, symmetric along the diagonal, where each cell is the classification accuracy (d’) for that pair of exemplars.

For each time point we compared each participant’s observed neural RDM with model RDMs, where each model RDM was a 120×120 matrix derived from theory, computational modelling, or behavioural data (as described in detail below). This analysis, known as ‘Representational Similarity Analysis’ (RSA) (Kriegeskorte, Mur, & Bandettini, 2008) tests the relationship between models of interest and the group data, measuring how well the model RDMs account for the observed pattern of results. At each time point we used Kendall’s tau-a to compute the rank order correlation between each candidate model and the neural data, and used these correlation values to compare candidate models in their ability to account for the neural data. Figure 2 shows the model RDMs, which are described in detail below.

**Figure 2.**
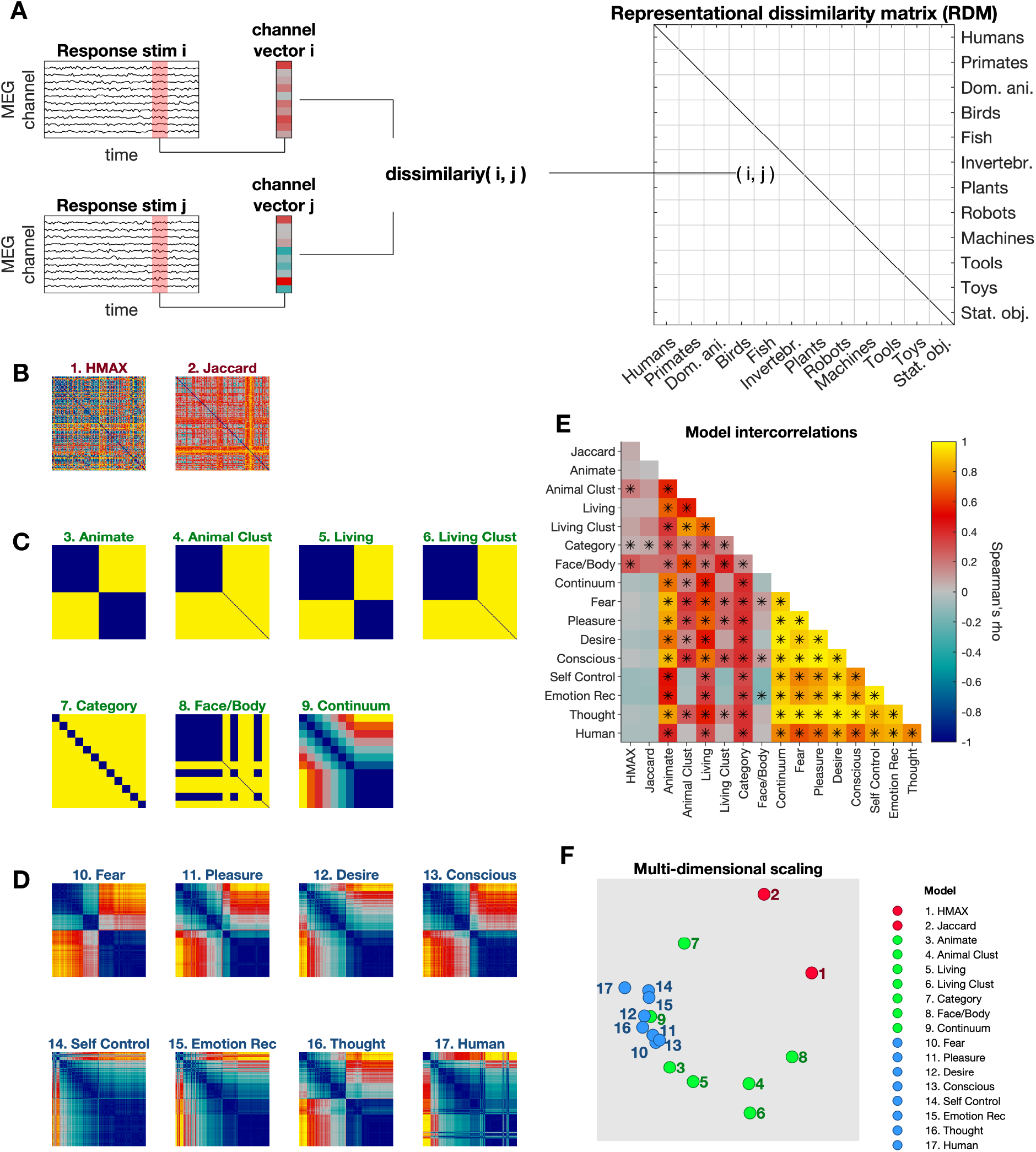
Models. *Representational Similarity Analysis (A).* For all pairs of images, the dissimilarity between their MEG response patterns is stored in a Representational Dissimilarity Matrix (RDM). An RDM is created for each time point from the MEG data. *RDMs used for model testing (B-D).* Model axes refer to all 120 image exemplars (grouped by category in the same order as Figure 2A. Colour bar indicates predicted degree of dissimilarity between exemplar pairs. Models are grouped according to whether they are low-level feature models (B), contemporary models (C), or behavioural models (D). *Model intercorrelations (E-F).* E) Model correlation matrix. Cell colour indicates correlation strength (\**p* < .05, adjusted for multiple comparisons across time points using a FDR of *q* < .01), with yellow cells indicating a stronger correlation between models, while blue indicates a weak/no correlation. F) MDS plot showing the representational geometry of model similarity in a 2 dimensional space. Models are colour coded according to whether they are low-level feature models, contemporary models, or behavioural models.

### Low-level feature models (Figure 2, models 1-2)

The HMAX and Jaccard silhouette models were included to test for the effects of low-level stimulus properties on the similarity/dissimilarity of neural responses, as measured using classifier performance.

#### HMAX (model 1)

Computational model of low-level visual processes. We applied the HMAX model (Riesenhuber & Poggio, 1999; Serre, et al., 2007) to simulate the responses of low-level visual areas. HMAX was applied to images at only a single image location and based on the standard orientation of each stimulus (i.e., not left-right flipped). The responses of the final HMAX layer (C2) for every stimulus were vectorized. We then generated the model RDM by taking the Euclidean distance between the vectorized model responses for each pair of stimuli.

#### Jaccard (silhouette model; model 2)

An abstract shape model that measures the shape of each object in terms of the pixels that the image occupies (Jaccard, 1901). We generated the model RDM by comparing the overlapping silhouette regions of two images at a time and obtaining a measure of the difference. This model was generated based on the standard orientation of each stimulus (i.e., not flipped), independent of location.

### Contemporary models of object representations (Figure 2, models 3-9)

The contemporary models were created based on organisational structures proposed in previous studies, with the term ‘contemporary’ used to highlight that these reflect current theories of object category structure. Descriptions of each model are provided below. *Dichotomy models (models 3 and 5):* The animate vs. inanimate dichotomy model (Caramazza & Shelton, 1998; Carlson, et al., 2013; Cichy, Pantazis, & Oliva, 2014; Kriegeskorte, Mur, Ruff, et al., 2008) is a category model that grouped all animate and inanimate objects separately (implying that objects within these groupings were more similar to each other, and more dissimilar to objects in the other grouping). Similarly, the living vs. non-living dichotomy model (Gainotti, 2000; Huth, Nishimoto, Vu, & Gallant, 2012; Warrington & Shallice, 1984) grouped all living and non-living objects separately. The living category included the same items as the animate category but with the addition of plants. *Cluster models (4 and 6):* The animal cluster model (model 4) is a single-category model that only grouped all animate objects together, suggesting that animate objects will be more similar to each other, and more dissimilar to all other objects, but that inanimate objects will not cluster. The living cluster model (model 6) follows the same principle, but grouping all living objects together. The cluster models were created to determine whether the effect of the dichotomy models was driven by cohesion within the in-group alone (i.e., animate, living), with more disparate object representations in the out-group category (i.e., inanimate, non-living) (Clarke & Tyler, 2014).

#### Faces/bodies model (model 7)

Faces and bodies stand out as special categories for object recognition (Barragan-Jason, Cauchoix, & Barbeau, 2015; Cauchoix, Barragan-Jason, Serre, & Barbeau, 2014; Gobbini, et al., 2011; Haxby, et al., 2001; van de Nieuwenhuijzen, et al., 2013) and so were of interest given the inclusion of toys and robots in our stimulus set. As such, the faces/bodies model is single-category model, grouping together all object categories that had faces or bodies, including all animate objects, as well as robots and toys.

#### Category model (model 8)

The category model was included as a measure of category individuation, as it proposes that items within individual categories have distinctly related patterns due to common visual and semantic properties, and these patterns are more different to those of objects from other categories (Clarke & Tyler, 2014). This model grouped each individual category as being more similar to within-category items and more dissimilar to other categories.

#### Continuum model (model 9)

The continuum model is a graded model based on the animacy continuum proposed by Sha et al. (2015). The continuum included a gradient of similarity between object categories that varied along a dimension related to biological classes, such that categories more similar to humans (biologically), would have more similar activity patterns, and those more dissimilar to humans would have activity patterns more similar to inanimate objects. For this model, plants were included on the continuum as they are a biological category and were represented on the continuum between invertebrates and inanimate objects. All non-living inanimate objects were treated as a single category, most dissimilar to the human category.

### Behavioural-rating models (Figure 2, models 10-17)

The behavioural-rating models include the agency/experience models (models 10 – 16) and the human model (model 17). These models were created by obtaining behavioural ratings of the stimuli according to a specific question (detailed below). A total of 325 Amazon’s Mechanical Turk workers residing in either the United States of America or Canada, completed one of the eight surveys online (number of participants per survey ranged from 40 – 43). Participants included 146 females (1 other, 1 no response), and had an average age of 35.27 years (SD = 10.26, range = 18.9 – 70.8; one age value missing). In each survey we asked workers to answer a single question for each of the stimuli:

*10. Fear* – How much is it capable of feeling afraid or fearful?

*11. Pleasure* – How much is it capable of experiencing physical or emotional pleasure?

*12. Desire* - How much is it capable of longing or hoping for things?

*13. Consciousness* - How much is it capable of having experiences and being aware of things?

*14. Thought* - How much is it capable of thinking?

*15. Emotion-recognition* - How much is it capable of understanding how others are feeling?

*16. Self-Control* - How much is it capable of exercising self-restraint over desires, emotions or impulses?

*17. Human* – How similar is this to a human?

Surveys for models 10-16 were based on a subset of the mental capacity surveys used in Gray et al. (2007), which vary as to how much they loaded onto the author’s ‘Experience’ and ‘Agency’ factors that were established in their study. The seven agency/experience models were based on the results of these surveys. The ‘Human’ survey (17) was added to address a meta-representational idea of categorization, that of “human-ness”: a complex factor which may encompass biology, agency, and visual similarity. Each survey required participants to rate all 120 images on a 7-point scale from ‘*Not at all’* to ‘*Very much so’* in response to the specific question. Each survey took approximately 10 minutes to complete and participants were financially compensated for their time. The surveys were created and administered using the Qualtrics online survey platform. For each survey, participants provided voluntary consent and basic demographic information before completing the survey. Participants were only allowed to complete one of the eight surveys available, resulting in unique individuals for each survey. Stimulus order was randomised separately for each participant.

To construct the models based on agency and experience (shown in Figure 2), an RDM was created for each set of survey responses by obtaining the absolute difference between image ratings for each pairwise comparison of the 120 images, using the mean ratings of each image. These RDMs, based on the survey ratings, provide hypothetical models of the degree of dissimilarity between the neural responses associated with each image. For graphical purposes, we scaled these difference values between 0 and 1 for each model, such that warmer colours indicate greater dissimilarity, while cooler colours depict greater similarity between the neural representations in the pair-wise comparison.

### Model intercorrelations (Figure 2E)

As the models we used in this study were not orthogonal, we measured the degree of overlap by performing correlations (Spearman) between each of the models (see Figure 2E). By evaluating the strength of these correlations, we obtained an estimate of how much the models overlap in terms of the hypotheses being tested. Of particular note, the behavioural-rating models based on the agency and experience factors from Gray, et al. (2007) and the human model we created were all highly correlated (see clustering in Figure 2F MDS plot of the representational geometry): this was not surprising as these models all capture slightly different aspects of similarity to humans.

In this study, we aimed to select stimuli that were visually diverse within each subcategory, to minimise the extent to which visual similarity would produce seemingly ‘categorical’ patterns of results. The model correlation data suggests that our stimulus set provided good separation of visual similarity and object category, since few models correlated with the visual feature models.

Importantly, this should minimise the contribution of low-level visual similarity when we evaluate our hypothesis driven models. Exceptions to this included the animal cluster, category, and faces/bodies models, which each showed a significant correlation with one, or both of the HMAX and Jaccard models. This suggests that despite our stimulus diversity, there was still greater visual homogeneity of exemplars within the category groupings in these models than between category groupings. This means that, particularly for the animal cluster, category, and faces/bodies models, any correlation between these models and the observed pattern of classifier performance could be driven by low-level visual similarity rather than by the higher-level category structure represented by these models.

## Results

### Decoding object exemplars from the MEG recordings

We scanned participants using MEG while they viewed 120 object stimuli and applied multivariate pattern analysis to the MEG sensor recordings at each time point, measuring how well the classifiers could decode the stimulus the participants were viewing. To study the brain’s representation of the objects at each time point, we ran the decoding analysis for all possible pairwise combinations of the 120 object stimuli. These data were used to create a set of time-varying RDMs identical in size to model RDMs (Figure 2).

We first confirmed we could decode the objects from the MEG recordings. Figure 3A shows the average performance of the classifier across all the pairwise combinations objects. The results show sustained decoding of object exemplars from 50 ms post stimulus-onset to the end of the time window (600ms) with peak decoding performance at approximately 105 ms post stimulus onset. These results are consistent with previous MEG decoding studies examining the emerging representation of objects in humans (Carlson, et al., 2013; Cichy, et al., 2014; Goddard, Carlson, Dermody, & Woolgar, 2016).

**Figure 3.**
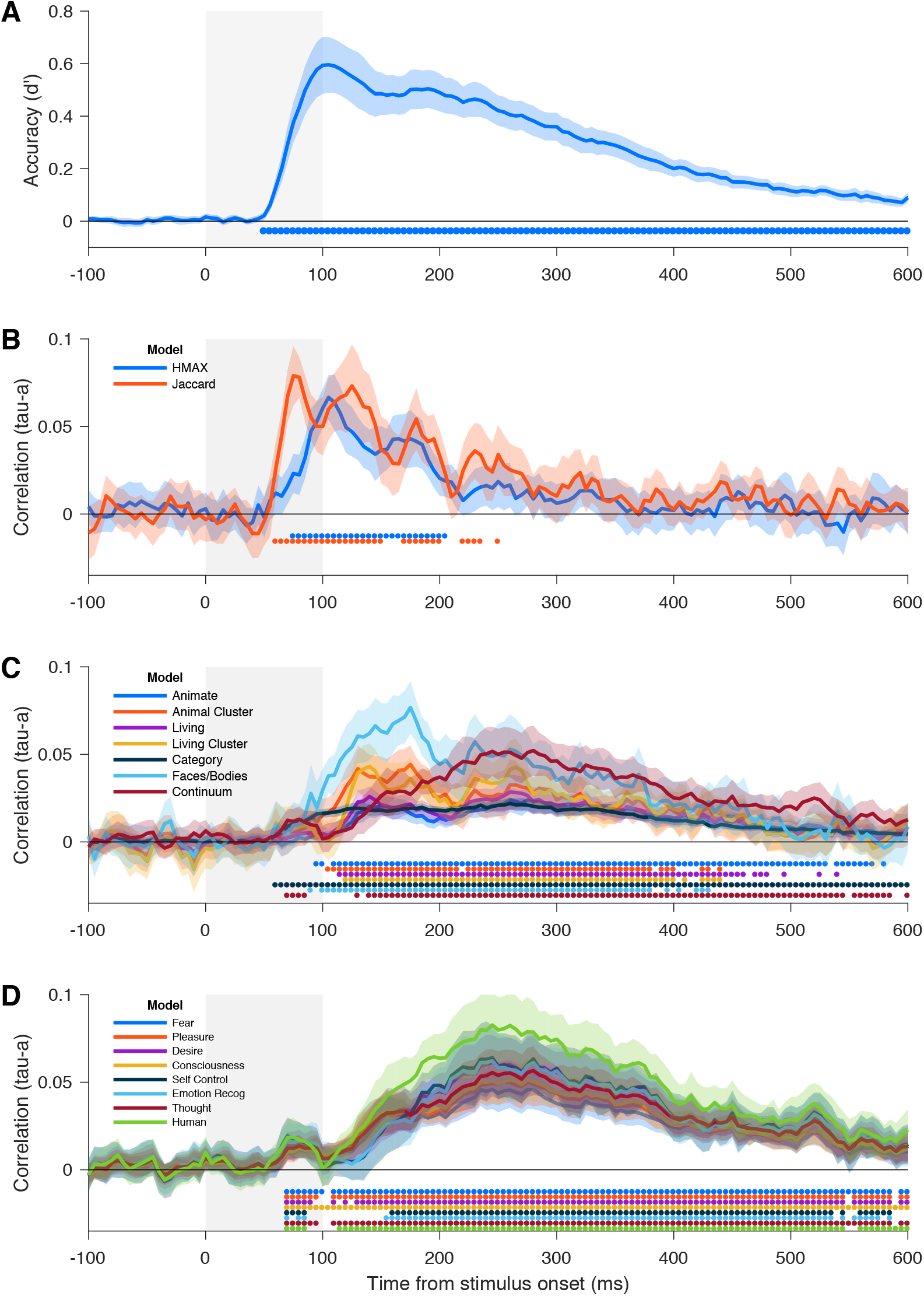
Exemplar decoding and model testing. ***(A) Exemplar decoding.*** Average decoding performance (measured in d-prime) over time for all exemplar pairs. Grey bar indicates the period the image was on the screen. Error bars indicate 95% between-subject confidence intervals. Blue dots along the x-axis indicate time points at which decoding performance was significantly above chance (one-tailed t-test, adjusted for multiple comparisons across time points using a FDR of *q* < .01). **(B-C) Model testing.** Correlation between the classifier data and (B) the low-level visual feature models, (C) contemporary models, and (D) behavioural-rating models. Grey bar indicates the period the image was on the screen. Shaded area indicates the 95% confidence interval of the between-subject means. Colour coded dots along the x-axis indicate time points where the model provides a significant account for the data using a Kendall’s Tau-a correlation (one-tailed t-test, adjusted for multiple comparisons across time points using a FDR of *q* < .01).

### The dynamic representation of objects

How does the brain’s representation of objects unfold over time? Having established that we could decode the individual object images, we next tested a range of hypotheses about category representations by comparing the observed neural RDMs with the model RDMs at each time point, using RSA (Kriegeskorte & Kievit, 2013; Kriegeskorte, Mur, & Bandettini, 2008). The neural RDM at each time point describes the brain’s representation of the stimuli at that time. The models (Figure 2B-D) attempt to explain a proportion of the variance in this structure. Formally, the models were evaluated by computing the rank correlation between the neural RDMs and each model RDM (Figure 3B-C).

### Low-level visual models account for early representations (Figure 3B)

We evaluated two low-level models to study how primitive visual features account for the brain’s representation of the stimuli. The Jaccard (i.e., silhouette) model evaluates the global shape of the stimuli (Jaccard, 1901). The HMAX model is based on a simulation of the response of early visual areas (Serre et al., 2007). Both the Jaccard and HMAX models were significantly correlated with the neural RDMs during early stages in the time course, peaking at 75 and 105 ms respectively, and were no longer significant predictors after 250 ms. This is in agreement with the established literature about the time-course of visual object recognition, with responses related to lower-level visual stimulus properties occurring earlier on, and more abstract semantic and categorical responses occurring later (Carlson, Simmons, Kriegeskorte, & Slevc, 2014; Carlson, et al., 2013; Cichy, et al., 2014; Clarke & Tyler, 2014). It should also be noted that the models show high correlations even though our design incorporated left-right flips of the images and spatial displacement of the images to reduce the influence of low-level stimulus properties. The low-level models were generated using only the standard orientation of each stimulus at a fixed position, yet could still predict the data after these transformations, affirming the importance of low-level visual similarity in the initial representation of the stimuli.

### Contemporary models: Intermediate processing emphasizes faces and bodies

A wide range of theoretical models have been proposed to account for the brain’s higher-order representation of objects. We tested how each of these models could account for the brain’s emerging representation of the objects (Figure 3B-D). The models we tested included a range of categorical models (e.g., animate versus inanimate), as well as a biological continuum model (Sha et al., 2015). We assessed their explanatory power using RSA and found that the models produced varying results. Starting at approximately 100ms, the face/body category model had the most explanatory power. Notably, the two other models with the early peaks (animal-cluster and faces/bodies) were among those showing significant overlap with one or both of the low-level feature models (see Figure 2E), suggesting that low-level visual similarity may have contributed to the earlier onset and high peaks for these models. At approximately 300ms, there appeared to be a transition in the representational structure. Here, the faces/body model, which was the best performing model in the early period (100-200ms), has declined and the biological continuum model increased its performance to have comparable explanatory power to the faces/bodies model. Interestingly, the animacy model was among the weakest performing models, despite a number of studies showing animacy provides a significant account of the human and primate brain’s higher-order representation of objects (e.g., Carlson et al., 2013; Cichy et al., 2014; Kiani et al., 2007; Kriegeskorte et al., 2008b).

### Human similarity model provides the best account of late representations

Higher order factors such as human similarity and agency are known to influence human perception of objects (Gobbini et al., 2011; Gray et al., 2007). To assess these attributes, we collected behavioural ratings for the stimuli about various higher order attributes (e.g., capacity to experience pleasure), to generate a new set of models. We then tested whether these models could account for the brain’s emerging representation of objects (Figure 3D). Across the behavioural-rating models, the results were very similar, which can be attributed to the high level of overlap in their internal structure (see Figure 2F). The models all show an initial peak at approximately 90ms, and then rise to a more significant peak at about 245 to 280 ms. This pattern was also observed for the biological continuum model, as seen in (Figure 3C). Notably, the best performing model in the later time window is the human similarity model, which is based on the question “How similar is this (object) to a human?”

### Human-ness and Agency/Experience models account for late representations

Our analysis of the models’ performance in the time series broadly indicated three distinct stages of processing. Early in the time series (<100ms) the low-level feature models performed the best. Next, in the intermediate time from 100ms to 200ms, three category models (Face/Body, Animal cluster, and Living Cluster) exhibited a distinct peak in their performance at about 180ms. Finally, the higher-order models based on agency and experience, and the biological continuum model, showed a slow rise that peaked about 270ms. To quantify these observations, we discretised the data in three 100ms time windows (0-100ms; 100-200ms; 200-300ms) in which we compared all model performances. Figure 4 shows the results of the windowed RSA analysis. Below each plot are visualisations of the representations at each stage constructed by projecting the data into two dimensions using t-SNE (Maaten & Hinton, 2008).

**Figure 4.**
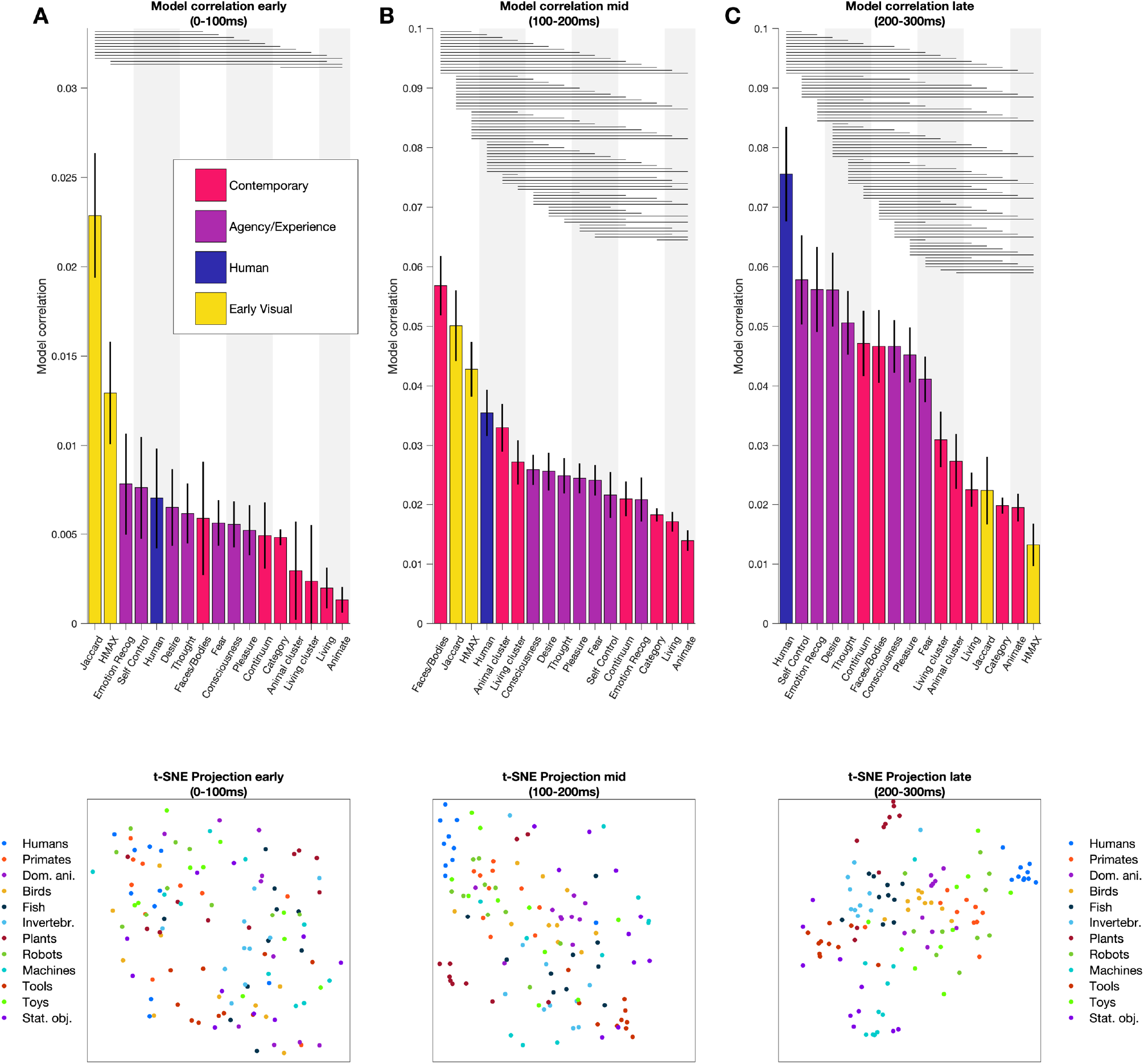
Model correlations (Kendall’s tau-a) in the early (0-100ms; A), middle (100-200ms; B) and late (200-300ms; B) time windows. Models are arranged in order of highest average rank correlation within each time window (highest on the left), and error bars indicate 95% SEM of the model correlation. Paired t-tests determined significant differences between model correlations (lines indicate significance at *p* < .05, adjusted for multiple comparisons across time points using a FDR of *q* < .01). Note the y-axis range for the early window is different because the model correlations were lower. Below each plot, two-dimensional embeddings of the stimuli are shown. The embeddings were computed using t-SNE (Maaten and Hinton, 2008). The distance between two points in this embedding reflects their neural dissimilarity.

To compare the models, we conducted a series of t-tests between all pairwise models to assess between-model performance in each of these time windows separately (adjusted for multiple comparisons across time points using an FDR of q < .01). In the early time window (0-100ms), the Jaccard (shape) model had the highest correlation (Figure 4A). In the second time window (100-200ms), the low-level models (Jaccard/ HMAX) still performed well; however, the face/body category model was the best performing model (Figure 4B). In the final time window (200-300ms), the low-level models (Jaccard/ HMAX) were among the worst performing models; and the human similarity model was the best model overall (Figure 4C). Notably, the humanness model outperformed the faces/bodies model, suggesting that the organisation of neural representations according to this human-centric model was not merely driven by the presence of faces/bodies, which are well-established as significant factors in object processing (Farah, 1996; Kanwisher et al., 1997; Thorpe et al., 1996). In terms of the model ranking, it can also be seen that the all the agency/experience models have good fits with the data, performing close or better than to the face/body category model and continuum model.

Overall, the results of the windowed analysis conforms with the observations from the time series data. Low level feature models provided the best fit for early representations, category models (in particular face/body, and the animal and living cluster) and low level feature models explained intermediate representations, and higher order models best explained late stage object representations. And, at this late stage of processing, the human similarity model provided the best fit to the data overall.

## Discussion

Many classifications, such as animate/inanimate and living/non-living, have been proposed as organisational principles for the brain’s representation of objects. Here we sought to provide an in-depth evaluation of contemporary models of visual object representations by evaluating their capacity to account for neural responses to a diverse range of object stimuli. In addition to these contemporary models and models based on low-level visual similarity, we created new theoretical and behaviour-based models. To test the predictive power of these models, we included novel stimuli that did not conform to the typical categories, such as robots and toys. Our results showed that the best performing model overall for late stage processing of objects was one based on the broad concept of human-similarity.

Our findings are consistent with accepted knowledge about the flow of information in human object recognition (for review see Contini, Wardle, & Carlson, 2017). This multi-stage processes begins with processing low-level visual properties of the stimulus, presumably in early visual cortex. These early representations are then subsequently transformed into higher order representations incorporating category structure (Carlson, et al., 2013; Cichy, Khosla, Pantazis, Torralba, & Oliva, 2016; Contini, et al., 2017) and semantic information (Carlson, et al., 2014; Clarke & Tyler, 2014). We found the best performing models at early time points were the low-level feature models (Jaccard and HMAX), and higher order models fit the data better later in the time series.

One of the most striking results was that one of the lowest performing models was the animate vs. inanimate model, despite being a well-established model in the literature (Caramazza & Shelton, 1998; Carlson, et al., 2013; Cichy, et al., 2014; Kiani, et al., 2007; Kriegeskorte, Mur, Ruff, et al., 2008; Proklova, Kaiser, & Peelen, 2016). Our study included stimuli that do not clearly have membership in the animate or inanimate categories (Bracci & Op de Beeck, 2016; Carlson, et al., 2013; Cichy, et al., 2014; Konkle & Caramazza, 2013; Kriegeskorte, Mur, Ruff, et al., 2008; Proklova, et al., 2016). The poor performance of the animate vs. inanimate model (and similarly the living vs. non-living model) likely could be accounted for by the inclusion of robots and toys. For example, visually inspecting the t-SNE plots (Figure 4) shows that robots are represented closer to humans and animate objects than to inanimate objects. This suggests that an animate/inanimate distinction is not the best way to classify these stimuli, and further highlights the impact of stimulus selection on defining the organisation of object categories (c.f. Carlson, Goddard, Kaplan, Klein, & Ritchie, 2018; Goddard, Klein, Solomon, Hogendoorn, & Carlson, 2018). Indeed, a recent fMRI study by Bracci, Kalfas, & Op de Beeck (2017) showed that visually confusing objects (e.g., a mug in the shape of a cow) exhibited neural activity patterns that were more similar to animate objects (i.e., an actual cow) than inanimates (Bracci, Kalfas, & Op de Beeck, 2017). Furthermore, as exemplar typicality affects the distinctiveness of category representations (Iordan, Greene, Beck, & Fei-Fei, 2016), the inclusion of these ambiguous object categories may have disproportionately affected a strict dichotomous categorisation model.

The superior performance of the human model builds on our existing understanding of the representation of object categories in the brain. The model extends on the continuum idea of Connolly et al. (2012) and Sha et al. (2015), as it represents a type of human-similarity continuum (see also Thorat, et al., 2019). However, unlike the animacy continuum that is based on biological classes, the human model was not limited by biology (Gobbini, et al., 2011; Tong, Nakayama, Moscovitch, Weinrib, & Kanwisher, 2000). Results from an fMRI study by Gobbini et al. (2011) are also consistent with a level of cross-over between animate/inanimate object categories that does not fit into this dichotomy, nor a continuum based on biological classes. The authors compared human observers’ perception of human faces and robots and found that robots evoked activation in areas associated with faces (though to a lesser extent than humans), while also activating object areas and areas associated with mechanical movements. This supports the idea of more a complex model of object categorisation that incorporates factors such as agency and human-related experiences. Given the relative strength of our human-centric model in accounting for the data, the idea of “humanness” as an important dimension in the neural representation of objects warrants further exploration.

Our best performing human-centric model likely encompasses a complex set of features, including both visual and conceptual factors. In our study, we did not impose a definition or any criteria against which people should rate the objects when asked ‘How similar is it to a human?’ (with responses from this survey used to generate the human model). Accordingly, we do not know which features people were using to rate object ‘humanness’, raising an interesting area for further investigation. The brain likely makes use of both visual and semantic information for representing objects (Carlson, et al., 2014; Clarke & Tyler, 2014; Coggan, Baker, & Andrews, 2016). Our data suggests that the semantic component of object representations incorporates information about concepts such as function, agency, and human experience. Indeed, a recent study by Connolly et al. (2016) showed an overlap between regions sensitive to the perceived threat of animals and those associated with social cognition, highlighting the importance of agent-related dimensions to object processing.

Presently, we still do not have a clear understanding of how different semantic concepts relate to object representations and category structure. A recent model attempts to explain the neural representation of object attempts using a multidimensional framework (Martin, 2016). In this paper, the author suggests that neural patterns associated with objects are formed from complex interactive circuits based on a range of systems throughout the brain, including those associated with action, perception and emotion. This idea shifts the focus away from models based on categories, with a view to a more holistic approach to object representations that considers interactions between various circuits throughout the brain. In this multidimensional framework, it is essential to recognize that no single feature or attribute could be able to fully explain the richness of the brain’s (multidimensional) representation of objects (see e.g., Thorat, et al., 2019). Recent fMRI studies have sought to identify principle axes of object representations in the brain (e.g., Connolly, et al., 2012; Sha, et al., 2015; Thorat, et al., 2019). In the present study, we show that the human similarity provided the best account of late stage processing, highlighting “humanness” as a key feature in the human brain’s representation of objects that shapes our experience of the world.

## Acknowledgements

This research was supported by an Australian Research Council Future Fellowship (FT120100816) and an Australian Research Council Discovery project (DP160101300) awarded to T.A.C. The authors acknowledge the University of Sydney HPC service for providing High Performance Computing resources. The authors declare no competing financial interests.

## Author contributions

E.C, E.G, M.W, T.C designed the study

E.C, T.G collected the data

E.C, E.G, T.G conducted the analysis

E.C, E.G, T.C interpreted the results

E.C and T.C wrote the manuscript

All authors reviewed the manuscript

